# Gene-resolved metagenomics reveals partitioned nitrogen transformation potential in a landfill leachate sequencing batch reactor microbiome

**DOI:** 10.64898/2026.11.09.763769

**Authors:** Samuel Rattray, Lynsay Blake, Nicola Ingrey, Iain MacArthur

## Abstract

Landfill leachate can contain high concentrations of ammonia and organic nitrogen requiring treatment before discharge. Sequencing batch reactors (SBRs) promote biological nitrogen removal through alternating aerobic and anoxic conditions, but taxonomic profiling alone cannot resolve how nitrogen transformation and competing retention pathways are distributed across microbial communities.

This study applied gene-resolved metagenomics to a sludge sample collected from a UK landfill leachate SBR. Nitrogen-metabolism genes were linked to contig-level taxonomic assignments and metagenome-assembled genome bins to examine nitrification, denitrification, dissimilatory nitrate reduction to ammonium (DNRA), and assimilatory nitrate reduction (ANR).

Nitrogen-transformation potential was partitioned across dominant and minor taxa. Nitrospirota and Pseudomonadota were the most abundant classified phyla, whereas Actinomycetota represented only 4% of the community but included Actinomycetes, which showed the highest within-taxon proportion of nitrogen metabolism associated genes. *amoABC* was assigned only to *Nitrosomonas* and Nitrososphaerota, whereas denitrification genes were more widely distributed but frequently occurred as partial pathways. *nosZ* was detected only in the minor phyla Chloroflexota, Acidobacteriota and Myxococcota. DNRA and ANR-associated genes occurred across multiple lineages; *nirA* in *Nitrospira* and DNRA-associated *nirBD* elsewhere within Nitrospirota suggested nitrifier-associated lineages may also encode nitrogen retention potential.

Gene-resolved metagenomics identified candidate low-abundance contributors, restricted distributions of genes associated with ammonia oxidation and nitrous oxide reduction, and nitrogen-retention potential not apparent from taxonomic composition alone. Because the study examined DNA-level potential in a single sludge sample, expression, reactor chemistry and longitudinal data are needed to establish pathway activity and operational relevance.

## Introduction

### Landfill leachate treatment and nitrogen pollution

An estimated 2.56 billion tonnes of municipal solid waste (MSW) were generated globally in 2022. Despite increasing emphasis on waste reduction and resource recovery, landfilling remains the most common waste management route globally [1].

During waste degradation, microbial decomposition mineralises organic nitrogen from solid waste proteins to ammonium, which can accumulate in the leachate [2]. Although ammonium can be oxidised through nitrification under aerobic conditions, landfill interiors rapidly become oxygen-depleted, favouring the persistence of ammonium while limiting nitrification and the production of nitrate and nitrite [3]. Denitrification may also be constrained in older landfills as readily biodegradable carbon becomes depleted and nitrate availability remains limited [3, 4].

Consequently, landfill leachate can contain high concentrations of soluble reactive nitrogen and requires treatment prior to discharge or transfer to downstream wastewater treatment [3, 5]. Where leachate or treated effluent enters receiving waters, elevated nitrogen concentrations can promote eutrophication, increasing oxygen demand and destabilising aquatic ecosystems [6-8]. Nitrite and nitrate contamination from landfills can also reduce drinking water quality and pose risks to human and environmental health [9]. Ammonia rich leachate can alter soil and groundwater chemistry, contributing to acidification and wider environmental stresses [10, 11].

Sequencing batch reactors (SBRs) are commonly used for biological treatment of nitrogen rich wastewater, including landfill leachate, to reduce ammonium and other nitrogenous contaminants to concentrations compatible with discharge requirements [12, 13]. SBR operation alternates between aerated and anoxic conditions, allowing multiple nitrogen transformation processes to occur within a single reactor. During aeration, nitrification converts ammonium to nitrate, while subsequent anoxic conditions promote nitrate reduction and denitrification. Treated wastewater is then separated from the settled sludge, while the microbial biomass is retained for subsequent treatment cycles [14-16].

### Nitrogen removal pathways in SBRs

The principal biological route for ammonium removal begins with nitrification. Ammonia oxidation is mediated by ammonia monooxygenase, encoded by *amoABC*. In ammonia-oxidising bacteria, subsequent hydroxylamine oxidation is associated with *hao*. Nitrite is then oxidised to nitrate by nitrite oxidoreductase, encoded by *nxrAB*. These reactions are frequently partitioned between ammonia-oxidising organisms such as *Nitrosomonas* and nitrite-oxidising organisms such as *Nitrospira* [17, 18]. Some *Nitrospira* can instead perform complete nitrification through the comammox pathway [19].

Under anoxic conditions, nitrate generated during nitrification can be removed from the system as gaseous N₂ through denitrification. Nitrate is initially reduced to nitrite by membrane associated nitrate reductases such as *narGHI* or the periplasmic *napAB* system. Nitrite is then reduced to nitric oxide through *nirK* or *nirS*, followed by nitric oxide reduction through systems including *norBC*. Finally, nitrous oxide is reduced to N₂ by nitrous oxide reductase, encoded by *nosZ* [20].

### Competing nitrogen retention pathways

Nitrate and nitrite within an SBR can also enter pathways that retain nitrogen within the system rather than removing it as nitrogen gas. DNRA competes directly with denitrification for nitrate and nitrite. The initial nitrate reduction step can utilise the same *nar* and *nap* systems as denitrification. DNRA subsequently reduces nitrite to ammonium through either the *nirBD* or *nrfAH* systems [21]. The resulting ammonium may therefore counteract the intended nitrogen removal function of the SBR. The factors controlling competition between denitrification and DNRA are not fully resolved, although DNRA can be favoured under high carbon-to-nitrate ratios and nitrate-limited conditions [22].

ANR provides an additional route of nitrogen retention. Rather than functioning primarily as respiratory nitrogen metabolism, ANR reduces nitrate and nitrite to ammonium for incorporation into cellular biomass. Nitrate reduction may involve assimilatory nitrate reductases encoded by *nasA/nasC* or *narB*, followed by assimilatory nitrite reduction through systems including *nirA* or *nasDE*. ANR is generally suppressed when ammonium is readily available because direct ammonium assimilation is energetically favourable [23]. Nevertheless, ANR genes have been detected in SBR communities [24], and assimilation into biomass represents a potentially important alternative fate for inorganic nitrogen [25]. Assimilation retains nitrogen within microbial biomass during reactor operation. However, biomass associated nitrogen can subsequently leave the treatment system through sludge wasting. Its contribution to overall nitrogen removal therefore depends on biomass retention, sludge withdrawal and the system boundary considered [26].

Consequently, nitrogen transformation within an SBR cannot be understood solely as nitrification followed by denitrification. Nitrate and nitrite occupy a central metabolic crossroad at which denitrification can remove nitrogen from the system, while DNRA can recycle it to ammonium and ANR can retain it within microbial biomass. Identifying organisms encoding complete or partial pathways associated with rate-limiting nitrogen-removal steps or competing nitrogen-retention pathways may reveal candidate functions relevant to nitrogen removal efficiency and the formation of undesirable products such as nitrous oxide.

**Figure 1.**
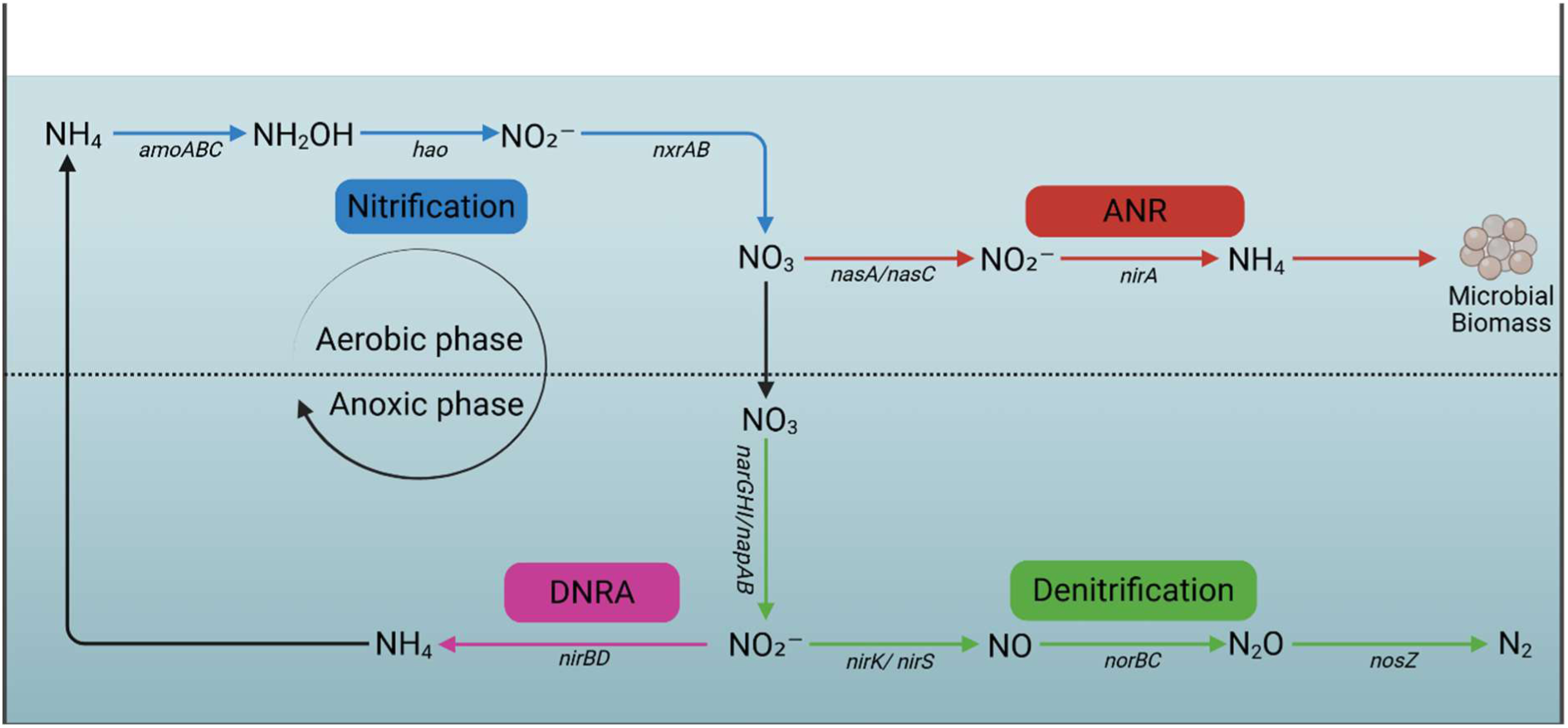
Nitrogen removal and nitrogen retention pathways provide alternative fates for nitrate and nitrite in SBRs. Genes shown are those identified in this study and do not represent an exhaustive set of genes involved in each pathway. During the aerobic phase, nitrification oxidises ammonium to nitrate through ammonia oxidation (*amoABC*, *hao*) and nitrite oxidation (*nxrAB*). During the anoxic phase, nitrate reduction (*narGHI/napAB*) supplies nitrite to denitrification, which produces nitrogen gas via nitric oxide and nitrous oxide (*nirK/nirS*, *norBC*, *nosZ*), or to dissimilatory nitrate reduction to ammonium (DNRA; *nirBD*), which returns nitrogen to the ammonium pool. Assimilatory nitrate reduction (ANR; *nasA/nasC*, *nirA*) converts nitrate to ammonium for incorporation into microbial biomass. Created with BioRender.com.

### Limitations in current understanding of SBR microbiomes

Previous studies of SBR microbial communities have frequently focused on changes in taxonomic composition associated with reactor operation, including organic loading, ammonium concentration, dissolved oxygen, alkalinity and wastewater type [17, 27-29]. They have identified that reactor performance can remain stable despite substantial changes in community composition because phylogenetically distinct organisms may occupy newly available niches and perform the same functional roles [27]. Despite this, functional interpretation of SBR microbiomes is often inferred from established ecology of taxa rather than through direct analysis of the genes responsible for individual nitrogen transformations [27, 30-40]. Consequently, functionally important low abundance taxa may remain unrecognised.

Where metagenomic approaches have been applied in SBRs and related treatment reactors, studies have examined laboratory scale mock-SBR rather than full scale industrial SBR systems [41-44], have not specifically resolved nitrogen metabolism genes [45-47], or have not investigated potentially competing pathways [41, 42]. Consequently, very few studies investigate how nitrogen removal and nitrogen retention pathways are genomically organised across microbial communities within full-scale landfill leachate SBRs [48]. In particular, the distribution and completeness of denitrification, DNRA and ANR pathways within individual taxa and reconstructed genomes remain poorly resolved.

### Study aim

This study applied gene resolved metagenomics to sludge from a full-scale landfill leachate SBR to examine how nitrogen-transformation potential was organised across the microbial community. Nitrogen metabolism genes were linked to contig-level taxonomic assignments and reconstructed genome bins to examine the distribution and completeness of nitrification, denitrification, DNRA and ANR pathways. The analysis assessed whether nitrogen-transformation potential was concentrated within dominant organisms encoding complete pathways or partitioned across multiple community members, and whether nitrogen-removal and nitrogen-retention potential coexisted within the microbiome.

## Methods

### Sample collection and processing

A single 1 L activated-sludge sample was collected on 15 October 2024 from a full-scale SBR treating landfill leachate in England, stored at 4 °C and processed within three weeks. The sludge was homogenised by mixing, and three 2 mL aliquots were withdrawn as technical extraction replicates. Each aliquot was mixed with 2 mL sterile 1× PBS and centrifuged at 10,000 × g for 10 min at 4 °C. The supernatant was discarded and the pellets were used for DNA extraction [49, 50].

### DNA extraction and purification

DNA was extracted using the FastDNA™ SPIN Kit for Soil (MP Biomedicals), including FastPrep™ mechanical lysis [49, 50], and purified using the genomic DNA cleanup protocol of the Monarch® Spin gDNA Extraction Kit (New England Biolabs). DNA was quantified using the Ǫubit™ dsDNA Broad Range Assay.

Purified DNA was assessed for amplifiability by PCR of the bacterial 16S rRNA gene using primers 27F/1492R and OneTaq® Ǫuick-Load DNA Polymerase. Cycling comprised initial denaturation at 94°C for 30 s; 30 cycles of 94°C for 20 s, 49°C for 30 s, and 68°C for 90 s; followed by a final extension at 68°C for 5 min and a hold at 4°C. A Milli-Ǫ water no-template control was included. Products were visualised on 1% agarose gels stained with SYBR™ Safe to assess amplifiability following purification. Purification optimisation, including comparisons with an alternative method, is shown in Supplementary Fig. S1.

### Nanopore library preparation and sequencing

DNA was normalised to 20 ng µL⁻¹ and libraries prepared using the ONT Rapid Barcoding Kit 24 V14 (SǪK-RBK114-24). Libraries were sequenced on a MinION Mk1D using an R10.4.1 FLO-MIN114 flow cell. Raw nanopore electrical signals were converted to nucleotide sequences (basecalled) in real time in MinKNOW v25.09.16 using Dorado v7.11.2 with the 400 bases s⁻¹ super-accurate (SUP) model; reads with Ǫ ≥10 were retained. Barcodes 1–3 yielded approximately 55130, 39550 and 69810 passing reads, comprising 263.12, 218.54 and 391.43 Mb, respectively. Read N50 values for barcodes 1–3 were 3,192, 5,100 and 3,185 bp, respectively.

### Metagenomic assembly and annotation

Metagenomic analysis was performed using SqueezeMeta v1.7.2 [51] in co-assembly mode. Reads were assembled using MEGAHIT v1.2.9 [52] with k-mer sizes 21, 29, 39, 59, 79, 99, 119 and 141, and contigs <200 bp were removed using PRINSEǪ [53]. rRNA and tRNA genes were identified using Barrnap and ARAGORN, respectively. Protein coding ORFs were predicted using Prodigal [54-57]. The resulting assembly comprised 102,981 contigs, with a total assembled length of 120,489,386 bp, and an N50 of 1,848 bp.

Genes were taxonomically annotated by DIAMOND searches against NCBI NR followed by SqueezeMeta lowest common ancestor assignment using the standard rank-specific identity thresholds, and functionally annotated against KEGG using the SqueezeMeta fun3 procedure [58, 59]. Both databases were supplied with the SqueezeMeta database bundle v1.8.0_20240607, built on 7 June 2024. Gene taxonomy was assigned to contigs using SqueezeMeta’s lowest common ancestor function. Reads from each replicate were mapped independently to the co-assembly using Minimap2 with the map-ont preset [60], from which gene TPM was retrieved.

### Taxonomic and nitrogen metabolism analysis

Taxonomic abundance was calculated as the proportion of all input reads mapping to contigs assigned to each taxon. Reads mapping to contigs without a resolved assignment at the relevant rank were retained as unclassified, while reads that did not map to the co-assembly were reported as unmapped. Unclassified assignments were retained in abundance denominators but excluded from analysis visualisations.

Downstream taxonomic, functional and abundance analyses were performed in R using custom scripts which are deposited publicly at samuel-rattray/SBR-nitrogen-metagenomics: R scripts for analysing taxonomic composition and nitrogen-transformation genes in a landfill leachate sequencing batch reactor (https://github.com/samuel-rattray/SBR-nitrogen-metagenomics/tree/main).

Candidate nitrogen transformation genes were identified from the KEGG gene annotation by gene name and KO numbers. Matches were classified as high confidence at ≥70% identity and query coverage and medium confidence at ≥40% identity and coverage; both required E-value ≤1 × 10⁻⁵ and alignment length ≥80 aa. KO confirmed matches below these thresholds were retained as low confidence, while unvalidated assignments were excluded.

Targeted genes comprised *amoABC, hao* and *nxrAB* for nitrification; *narGHI, napAB, nirK, nirS, norBC* and *nosZ* for denitrification; *nar/nap, nirBD* and *nrfAH* for DNRA; and *narB, NR, nasA/nasC, nasB, nirA, NIT-C, nasD* and *nasE* for ANR, *hdh* and *hzsA* for anammox.

Functional abundance was normalised within each taxon following the taxon-specific TPM rescaling approach implemented in SǪMtools [61]. For each taxon, the within-taxon relative abundance of nitrogen-metabolism-associated genes was calculated as:

> Target gene enrichment within taxon (%) = 100 × Σ target gene TPM / Σ total gene TPM assigned to taxon.

Because the data were derived from metagenomic DNA, TPM here describes mapped DNA abundance rather than transcription. For pathway heatmaps, TPM values for each gene were first averaged across the three extraction replicates. For each genus or phylum, the mean gene TPM values were summed and divided by the summed mean TPM of all genes assigned to the same taxon.

### Metagenomic binning and genome-bin analysis

Bins were reconstructed from the co-assembly using CONCOCT v1.1.0 [62] and MetaBAT2 [63] using sequence composition and differential coverage across the three replicates. CONCOCT used 10-kb contig fragments for initial clustering. Bins were integrated using DAS Tool [64], and taxonomy was assigned by SqueezeMeta from constituent contigs.

Completeness and contamination were estimated using CheckM2 v1.0.2 [65]. Bins with ≥30% completeness and ≤10% contamination were retained for gene presence analysis only. Bins below the 50% completeness were treated as partial bins. Bins without a resolved taxonomic assignment were retained as unclassified.

Retained bins were screened for the same nitrogen transformation genes and validated using the same procedure described above. Mean gene TPM across the three extraction replicates was normalised as:

> Bin normalised gene abundance = gene TPM / total Bin TPM.

**Table 1.** Characteristics and detected nitrogen-transformation genes of the fourteen retained genome bins. Combined gene labels indicate unresolved functional assignments. Missing markers do not establish biological absence.

| **Bin ID** | **Size (Mb)** | **Completeness (%)** | **Contamination (%)** | **Taxonomy** | **Detected nitrogen-transformation genes** |
|---|---|---|---|---|---|
| Nitrospiraceae_13 | 4.306 | 80.69 | 3.60 | Nitrospirota; Nitrospiraceae | *napA, narG/nxrA, narH/nxrB, nirA, nirD, nirK, nirS* |
| unclassified_12 | 5.961 | 51.30 | 3.23 | Unclassified Bacteria | *narG/nxrA, narH/nxrB, nirB, norB, nrfA* |
| Chloroflexota_7 | 3.199 | 47.77 | 0.22 | Chloroflexota | *narG/nxrA, narH/nxrB, nirD, nirK, nosZ* |
| unclassified_44 | 4.974 | 43.53 | 1.24 | Unclassified Bacteria | *narG/nxrA, narH/nxrB, nirD, norB, norC, nosZ* |
| unclassified_33 | 3.712 | 39.89 | 2.96 | Unclassified Bacteria | *amoA, amoB, amoC, nasA/nasC, nirB, nirK, norB, nrfH* |
| unclassified_58 | 3.930 | 39.30 | 3.44 | Unclassified Bacteria | *napA, narG/nxrA, narH/nxrB, nirB, nirS, nosZ, nrfA* |
| unclassified_43 | 3.797 | 38.54 | 5.24 | Unclassified Bacteria | *narI, nirB, nirD* |
| Nitrospirales_5 | 2.357 | 38.29 | 1.58 | Nitrospirota; Nitrospirales | *napA, nirD, nirK, norC* |
| unclassified_2 | 4.740 | 36.07 | 4.46 | Unclassified Bacteria | *narG/nxrA, narH/nxrB, nasA/nasC, nirK, norB, nosZ* |
| unclassified_7 | 2.929 | 34.97 | 2.48 | Unclassified Bacteria | *amoB, napA, narG/nxrA, narH/nxrB, narI, nasA/nasC, nirB, nirK, nirS, norB* |
| unclassified_51 | 3.785 | 33.54 | 5.69 | Unclassified Bacteria | *napA, narH/nxrB, narI, nasA/nasC, nirA, nirK* |
| Nitrospirales_27 | 3.073 | 33.37 | 1.61 | Nitrospirota; Nitrospirales | *napA, narG/nxrA, nasA/nasC, nirA, nirD, nirK, norB* |
| unclassified_29 | 2.700 | 32.87 | 3.58 | Unclassified Bacteria | *narH/nxrB, nasA/nasC, nirB, nirD* |
| unclassified_19 | 3.235 | 30.01 | 1.30 | Unclassified Bacteria | *narG/nxrA, narH/nxrB, nasA/nasC, nirS* |

## Results

### Bin reconstruction of nitrogen pathway organisation

Genome bins were reconstructed to examine gene co-occurrence within the co-assembly [48]. Because the dataset comprised only three extraction replicates of the same sludge sample, the differential coverage information available for genome binning was limited. Fourteen bins met the retention criteria: three assigned to Nitrospirota, one to Chloroflexota and ten without a resolved taxonomic assignment. These bins support the taxonomic assignments and gene co-occurrence results described below and are fully described in Supplementary Fig. S2.

### Taxonomic abundance does not capture all nitrogen-transformation potential

To assess whether taxonomic abundance scales with nitrogen transformation potential, the ten most abundant microbial classes were compared for community abundance and their proportion of validated nitrogen metabolism associated genes (Fig. 2). Of the total input read abundance, 44% could be assigned to the bacterial kingdom, 0.2% to archaea, and 0.009% to eukaryotes. 51% of reads were not incorporated into the assembly and were therefore classified as unmapped. A further 4.67% of reads mapped back to the assembly but lacked a taxonomic match and were classified as unclassified, representing poorly characterised microorganisms. Because unclassified annotation can group multiple unrelated microorganisms, especially at lower taxonomic levels, unclassified taxa were excluded from this analysis but included in the community abundance calculations to avoid artificially inflating classified groups.

At broad taxonomic levels, the community contained groups previously reported in landfill leachate SBR microbiomes, including Pseudomonadota, Actinomycetota, and Nitrospirota [29, 30, 48]. Planctomycetota, Candidatus Sumerlaeota and Nitrososphaerota were also detected. Bacterial taxonomic composition is shown in Supplementary Fig. S3, and the full interactive taxonomic breakdown is provided in Supplementary File S1.

Actinomycetota accounted for 4% of total community abundance and was represented by the classes Actinomycetes, Thermoleophilia and Acidimicrobiia. These classes have previously been reported as minor members of SBR and wastewater bioreactor communities [66-68]. Actinomycetes had the highest proportion of nitrogen transformation genes of all identified classes (1.40%), suggesting disproportionate nitrogen transformation potential relative to their abundance. This was notable because, although Actinomycetota have established denitrification and DNRA capabilities [69], they are not typically regarded as core SBR nitrogen-removal taxa [27, 30, 66]. Acidimicrobiia and Thermoleophilia showed lower proportions of nitrogen-metabolism associated genes (0.39% and 0.05%, respectively), consistent with previous associations of Acidimicrobiia with denitrification and DNRA [70] and Thermoleophilia with ANR [71]. A higher within taxon abundance does not establish the greatest contribution to community-wide nitrogen transformation, but it highlights Actinomycetota as a potentially under recognised reservoir of nitrate and nitrite transformation potential in landfill leachate SBR sludge.

Nitrospirota represented 12.33% of the total community abundance, of which 9.67% was resolved to the class Nitrospiria. Nitrogen metabolism associated genes represented 0.74% of ORFs assigned to Nitrospiria.

The second most abundant phylum was Pseudomonadota (9.00% of total community abundance), represented by the classes Alphaproteobacteria (2.67%), Betaproteobacteria (4.67%), and Gammaproteobacteria (0.57%). All three classes showed similar levels of nitrogen metabolism associated gene abundance (0.27-0.55%). Members of Pseudomonadota are known for their metabolic versatility, and all three classes have been associated with both nitrification and/or denitrification [72-75]. These findings indicate nitrogen transformation potential across multiple classes of Pseudomonadota.

The phylum Planctomycetota accounted for 0.93% of the community. Within this phylum, Planctomycetia accounted for 0.23% of the reported community abundance, but showed no nitrogen metabolism genes. Despite Planctomycetota including branching anammox related lineages [76], no anammox genes were detected in this dataset.

Although Archaea represented only a small fraction of the community, the phylum Nitrososphaerota accounted for most archaeal abundance (0.13%). This phylum was exclusively represented by the class Nitrososphaeria, which showed a measurable level of nitrogen metabolism associated genes (0.78%) within this class. Members of Nitrososphaeria are well established ammonia oxidising archaea (AOA) [77]. AOA have been reported in SBR environments and are often outcompeted by AOB due to their greater growth rates, however AOA’s higher affinity to ammonia means it can outcompete AOB in ammonia deficient conditions [78, 79].

The candidate phylum Candidatus Sumerlaeota accounted for 1.53% of the reported community abundance and was represented by a single candidate class. No nitrogen transformation genes were identified in this phylum. Although *Candidatus* Sumerlaeota has been suggested to engage in DNRA [80] and detected in environments similar to SBR systems [81], genes associated with these functions were not detected in this dataset.

Nitrogen metabolism associated genes were distributed across multiple taxa but did not scale directly with abundance. Dominant groups such as *Nitrospiria* and Pseudomonadota classes showed measurable within taxon proportions of nitrogen metabolism associated genes, yet lower abundance taxa, particularly Actinomycetes, exhibited a disproportionately high proportion of nitrogen metabolism associated genes relative to their community abundance.

**Figure 2.**
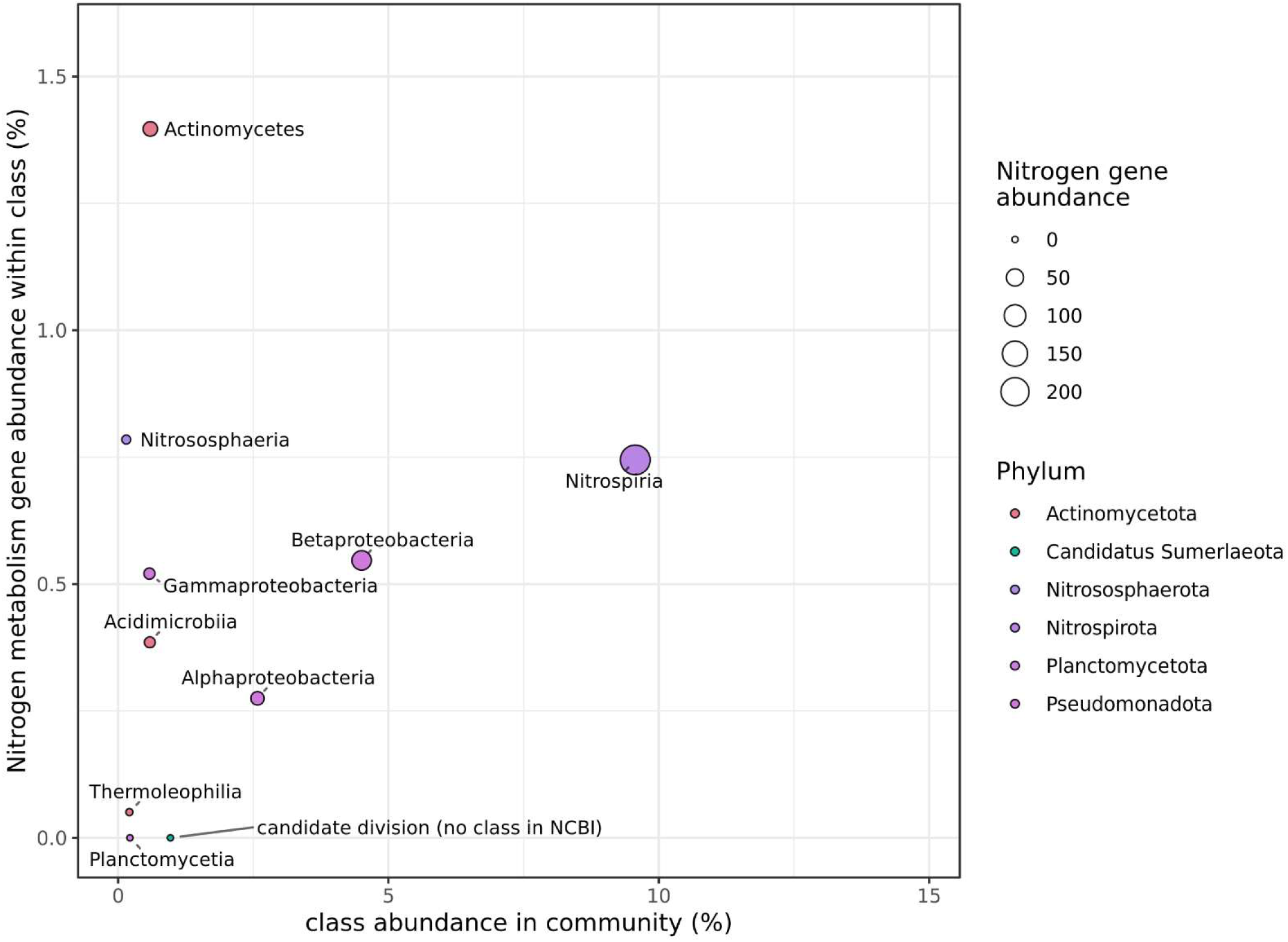
Actinomycetes show the highest within-taxon relative abundance of nitrogen-metabolism genes despite their low community abundance. Scatter plot showing mean class level community abundance across three extraction replicates against the proportion of nitrogen metabolism associated genes, calculated as the summed TPM of validated nitrogen metabolism genes divided by the total gene TPM assigned to that class × 100. Each point represents a microbial class, with point size indicating summed validated nitrogen gene abundance and fill colour indicating parent phylum. Unclassified taxa were retained in abundance calculations but excluded from the visualisation.

### Ammonia oxidation genes are taxonomically restricted, while nitrite oxidation genes are more widely distributed

In SBRs, nitrification is thought to be conducted by *Nitrosomonas*, oxidising ammonia and hydroxylamine, and *Nitrospira,* which oxidises nitrite [27, 30, 35, 36, 39, 82, 83]. As expected then, *amoABC* was associated with *Nitrosomonas*, and *nxrAB* associated with *Nitrospira*. However additional taxa carrying nitrification genes were identified within the dataset.

Among classified taxa*, amoABC was assigned to Nitrosomonas* and Nitrososphaerota, indicating a potential archaeal contribution to ammonia oxidation in this landfill leachate SBR. This may be particularly relevant for SBRs under conditions that favour archaeal ammonia oxidisers [77, 84]. For both *Nitrosomonas* and Nitrososphaerota, all three subunits of the *amoABC* complex were detected, indicating genetic potential, but also a taxonomically restricted step of nitrification in the system. All three *amoABC* subunits were also co-detected only in the unclassified genome bin unclassified_33.

Additionally, *hao*, which is associated with subsequent oxidation of hydroxylamine to nitrite, was assigned only to *Nitrosomonas.* Laboratory-scale SBR studies have shown that disturbances affecting *Nitrosomonas* can substantially inhibit ammonia oxidation [85]. The restricted distribution observed here suggests that loss of *Nitrosomonas* could constrain ammonia oxidation and downstream nitrogen transformations, while Nitrososphaerota may provide alternative ammonia-oxidation potential.

*nxrA* and/or *nxrB* were detected in Pseudomonadota, Planctomycetota, Chloroflexota, Acidobacteriota, Verrucomicrobiota, Nitrospirota with *Nitrospira*, and Actinomycetota, with *Mycobacterium* and *Mycolicibacterium*. *Nitrospira* is considered the principal nitrite oxidiser in SBR [17, 18], although Pseudomonadota [66, 86] and Chloroflexota [87] have also been reported as dominant nitrite oxidisers in SBR systems. Environmental *Mycolicibacterium* has previously been identified with *nxrAB* through culturing [88]. Therefore, SBR nitrite oxidation may not be entirely reliant on *Nitrospira* but may involve a wider community. Further high resolution metagenomic studies could identify other SBR nitrite oxidisers.

Because *narGH* and *nxrAB* are highly homologous some *nxrAB* assignments may instead be *narGH*. However, this broad taxonomic distribution of *nxrAB* has been reported in non-SBR aquatic environments, using approaches with enough resolution to differentiate *nxrAB* and *narGH* [89, 90]. These observations provide some support for the annotations, although the functions cannot be distinguished from sequence similarity alone.

**Figure 3.**
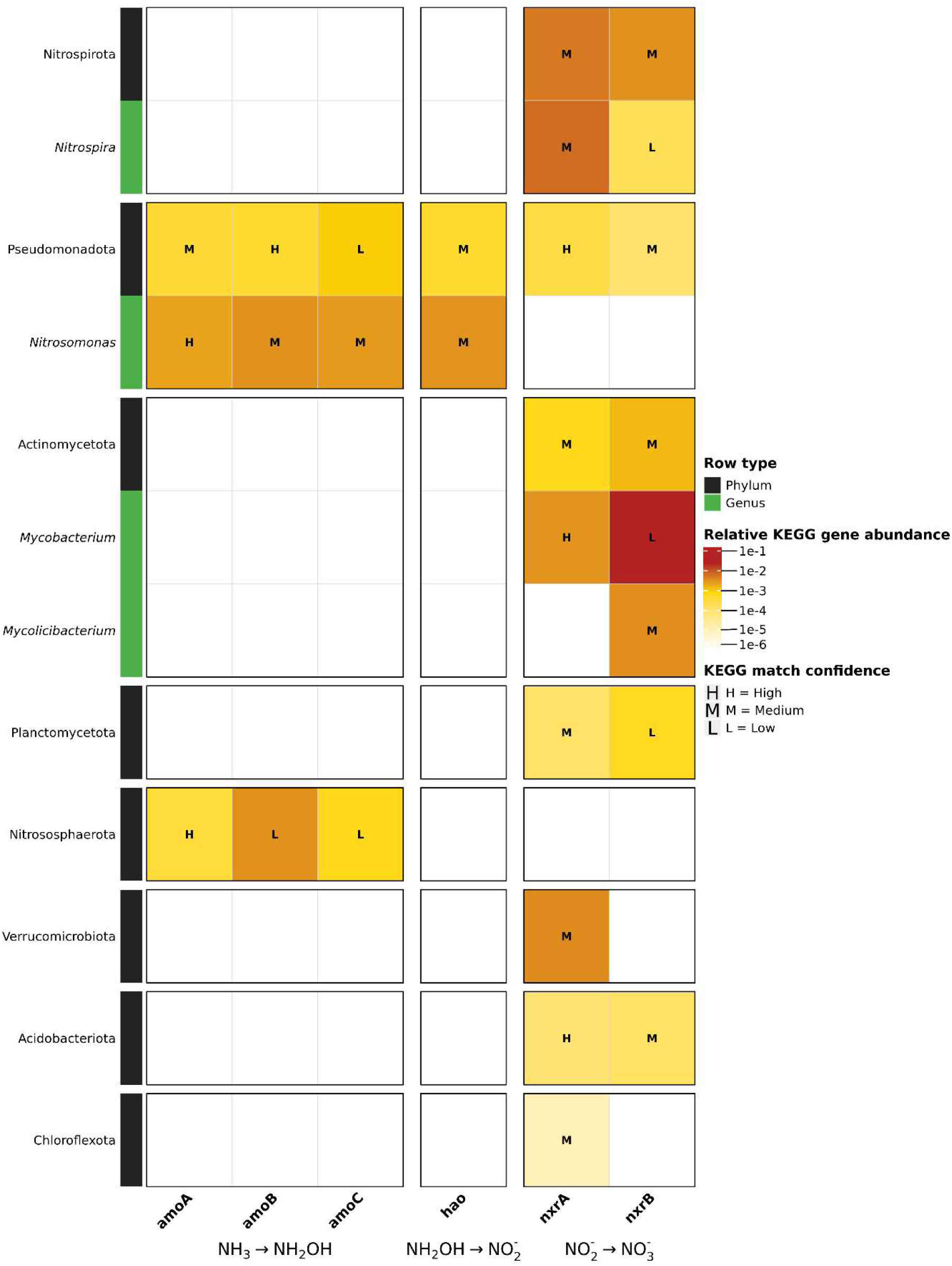
Detected ammonia oxidation genes are restricted to *Nitrosomonas* and Nitrososphaerota. Heatmap showing the distribution of nitrification associated genes across classified phyla and genera. Rows are grouped by phylum and include phylum level summaries and resolved genus level assignments. Columns represent genes involved in ammonia oxidation to hydroxylamine, hydroxylamine oxidation to nitrite and nitrite oxidation to nitrate. Colour intensity represents log10-normalised gene abundance, calculated from mean TPM across the three extractions.

### Denitrification genes are widely distributed, but nitrous oxide reduction potential is restricted to minor taxa

Denitrification genes were broadly distributed, suggesting that denitrification potential was partitioned across different community members [91]. *narGHI* genes were found across Nitrospirota, Pseudomonadota, Actinomycetota, Chloroflexota, Acidobacteriota, Planctomycetota, Verrucomicrobiota and were assigned at genus level in *Nitrospira*, *Stutzerimonas*, *Mycobacterium* and *Mycolicibacterium*. In contrast, *napA* was detected only in Nitrospirota and Pseudomonadota, including *Novosphingobium.* Thus, *narGHI* was the more widely detected nitrate reduction system in the dataset.

Nitrate reduction by *Nitrospira* has been reported in SBR like systems [92], while *Mycobacterium* and *Mycolicibacterium* are known to encode *narGHI* in other contexts [93, 94]. Among the *nar* genes, only *narG* and *narH* were present with *narI* absent, except in *Stutzerimonas*, where only *narI* was detected. Bin reconstruction reinforced this: Nitrospiraceae_13 encoded *narGH*, Nitrospirales_27 encoded only *narG*. Only in unclassified_7 were all three *narGHI* genes identified, suggesting that a full *narGHI* complex may be restricted to poorly classified genomes in this dataset. However, *narGH* nitrate reduction can occur without *narI* under some electron donor conditions [95], meaning *narGH* detection may still represent nitrate reduction potential.

The less widely distributed *napAB* system showed a similar pattern of incomplete gene recovery. *napA* was detected in Nitrospirota and Pseudomonadota with *Novosphingobium* which has established nitrate reducing potential [96], although the role of *napA* in SBR associated taxa, remains unclear.

Some Nitrospirota are known to encode *napAB* [97]. Because *napB* is required for canonical NapAB mediated nitrate reduction [98], the isolated *napA* detection may indicate incomplete pathway recovery or non-canonical nitrate reduction. Nitrospirota strains under SBR-like conditions have been reported to transcribe *napA* but not *napB* [99], indicating *napA* expression can occur without *napB*, although the exact function remains unclear.

Nitrite reduction to nitric oxide has two distinct routes using either *nirK* or *nirS*. Both genes were detected in Pseudomonadota and Nitrospirota. A single *nirS* signal was assigned to *Nitrospira* and was also recovered from the bin Nitrospiraceae_13. However existing studies that report *Nitrospira* encodes *nirK* rather than *nirS* in other SBR and activated sludge environments [100, 101]. *nirK* was detected in Nitrospirota and across all Nitrospirota bins. Together, these data support *nirK* associated nitrite reduction potential within Nitrospirota, but the assignment of *nirS* to *Nitrospira* requires further investigation.

*nirK* was more widely distributed, also found across Gemmatimonadota, Nitrososphaerota, Chloroflexota, Actinomycetota, and several members of Pseudomonadota, including *Permianibacter* and *Nitrosomonas.* Supported by bin analysis where *nirK* was detected within *Chloroffexota_7,* and multiple unclassified bins. The broader distribution of *nirK* is consistent with observations from other environments, where *nirK* often occurs across a wider taxonomic range than *nirS* [102]. The detection of *nirK* in *Nitrosomonas* and *Nitrosarchaeum* is consistent with previous SBR evidence [48, 103], whereas its assignment to *Permianibacter* has not been corroborated in the available literature and requires further validation.

Nitric oxide reduction showed a more restricted distribution. Both *norB* and *norC* were detected in Pseudomonadota, including *Nitrosomonas* consistent with previous SBR studies [103]. In Campylobacterota, only *norB* was detected, consistent with other SBR studies [104]. Nitric oxide reduction may instead involve qNOR, a single subunit NorB-like enzyme that uses quinol as the electron donor and does not require NorC [105].

The final denitrification step showed the most restricted gene distribution. *nosZ*, was detected only in Chloroflexota, Acidobacteriota and Myxococcota. Bin analysis identified Chloroflexota_7 plus three unclassified bins to have *nosZ*. This distribution is consistent with wastewater treatment studies, though not specifically SBR studies, that have reported *nosZ* containing Chloroflexota [106], Acidobacteriota [107] and Myxococcota [108]. This indicates strong genetic potential for nitrous oxide reduction but also highlights how *nosZ* is only identified in minor and poorly understood taxa in the system.

**Figure 4.**
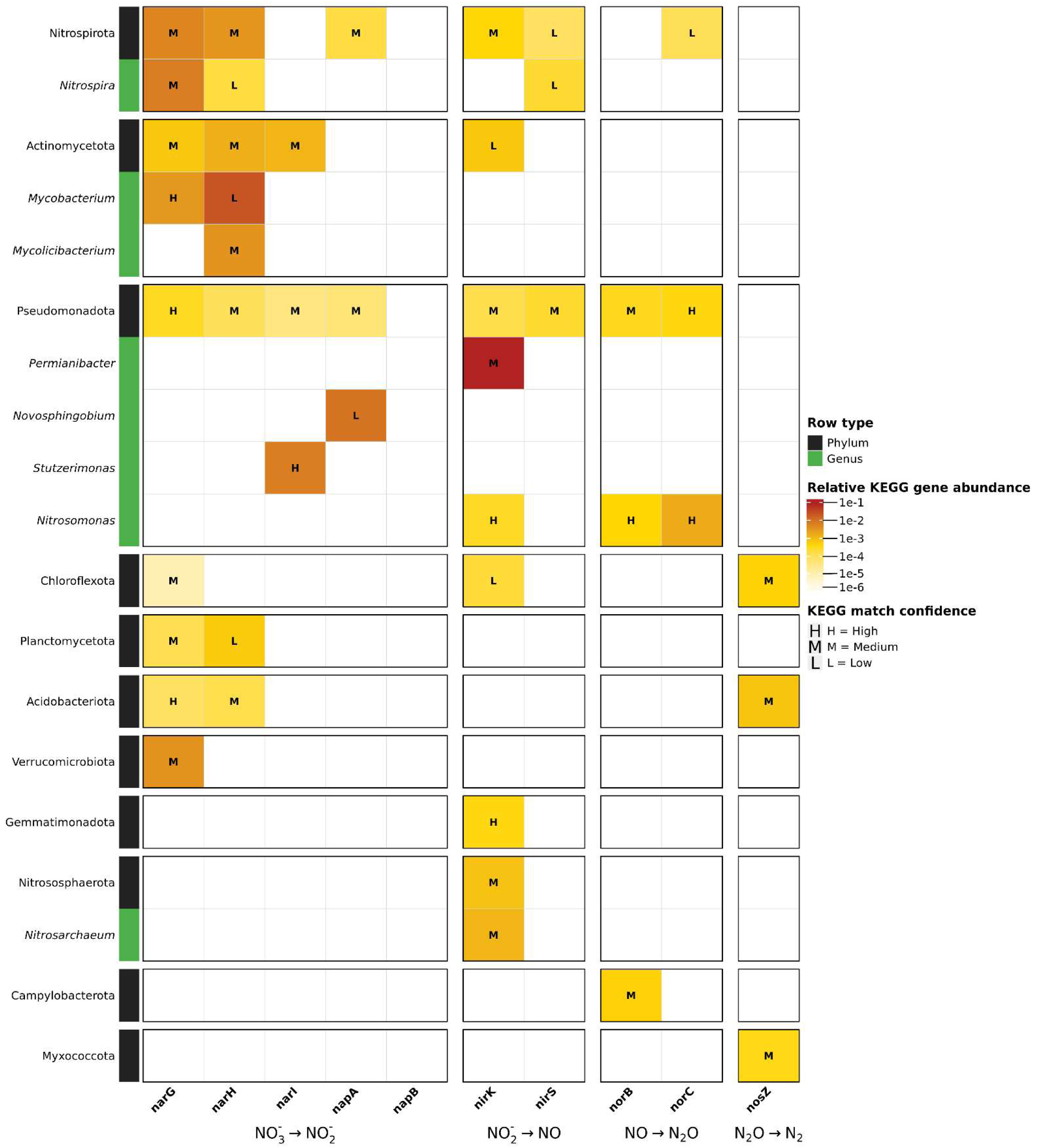
Denitrification genes are widely distributed, but detected nitrous oxide reduction potential is restricted to minor taxa. Heatmap showing the distribution of denitrification associated genes across classified phyla and genera. Rows are grouped by phylum and include phylum level summaries and resolved genus level assignments. Columns represent genes associated with nitrate reduction, nitrite reduction, nitric oxide reduction and nitrous oxide reduction. Colour intensity represents log10-normalised gene abundance, calculated from mean TPM across the three extraction replicates.

### Nitrogen-retention potential occurs across multiple lineages, including taxa associated with nitrification

ANR has previously been proposed in high bioload SBRs [109]. In this study, the nitrate reductase genes *nasA*/*nasC* were detected across Chloroflexota, Omnitrophota, Actinomycetota and Pseudomonadota, including *Pseudomonas*, whereas *nasB* was restricted to Pseudomonadota. Because both components are normally required for assimilatory nitrate reduction [110], the combined detection of *nasA*/*nasC* and *nasB* provides the strongest evidence for ANR potential within Pseudomonadota, consistent with previous SBR studies [111, 112]. In contrast, isolated *nasA*/*nasC* detections in the other phyla provide only limited evidence and may reflect incomplete pathway recovery or alternative gene functions, as similar detections without *nasB* have been reported in other SBR metagenomes [113, 114].

Genes associated with ANR nitrite reduction were even more restricted. Only *nirA* was detected among genes associated with ANR nitrite reduction. Its assignment to *Nitrospira* was supported by detection in the genome bins Nitrospiraceae_13 and Nitrospirales_27. *nirA* has previously been detected in SBRs [115] and identified as part of *Nitrospira* genomes in wastewater treatment facilities [116]. These findings indicate that *Nitrospira* may encode an under-recognised ANR potential in SBR systems, although the presence of *nirA* alone does not indicate a complete ANR pathway.

DNRA associated genes have previously been detected in SBR samples [21, 22], with DNRA activity demonstrated alongside denitrification as part of standard SBR operation [117]. In this study, putative DNRA potential was associated primarily with *nirBD* rather than *nrfAH*. Only *nrfA* was detected from the *nrfAH* system among classified taxa. This assignment was low confidence and restricted to Pseudomonadota. In contrast, *nirBD* genes were widespread, consistent with previous SBR and SBR-like metagenomic studies that reported greater *nirBD* than *nrfAH* abundance [44, 115, 118, 119]. *nirB* and/or *nirD* were detected in Nitrospirota, Bacteroidota, Acidobacteriota, Planctomycetota, Actinomycetota, including *Mycobacterium*, and Pseudomonadota, including *Mesorhizobium*. Most of these assignments cannot be corroborated at phylum level; however DNRA-focused SBR studies have identified Bacteroidota and Pseudomonadota as DNRA bacteria [117, 120].

Although Pseudomonadota are established contributors to DNRA in SBRs [121], *Mesorhizobium* has not previously been implicated in this setting. Its assignment here is nevertheless plausible because *nirBD* has been reported in *Mesorhizobium* genomes from other environments [122]. Similarly, *Mycobacterium* has demonstrated DNRA potential outside SBR systems [123]. These findings therefore identify *Mesorhizobium* and *Mycobacterium* as candidate carriers of DNRA associated genes, although their DNRA activity in SBRs requires further validation.

Nitrospirota are strongly associated with nitrification, as discussed above. Detection of *nirBD* within this phylum is therefore notable because it suggests that lineages associated with nitrification may also encode nitrogen retention potential. Although DNRA genes have not been widely reported in Nitrospirota from SBRs, they have been identified in Nitrospirota from other environments [97]. Together with the detection of *nirA* in *Nitrospira*, these findings suggest that members of Nitrospirota may encode functions associated with both nitrification and nitrogen retention.

**Figure 5.**
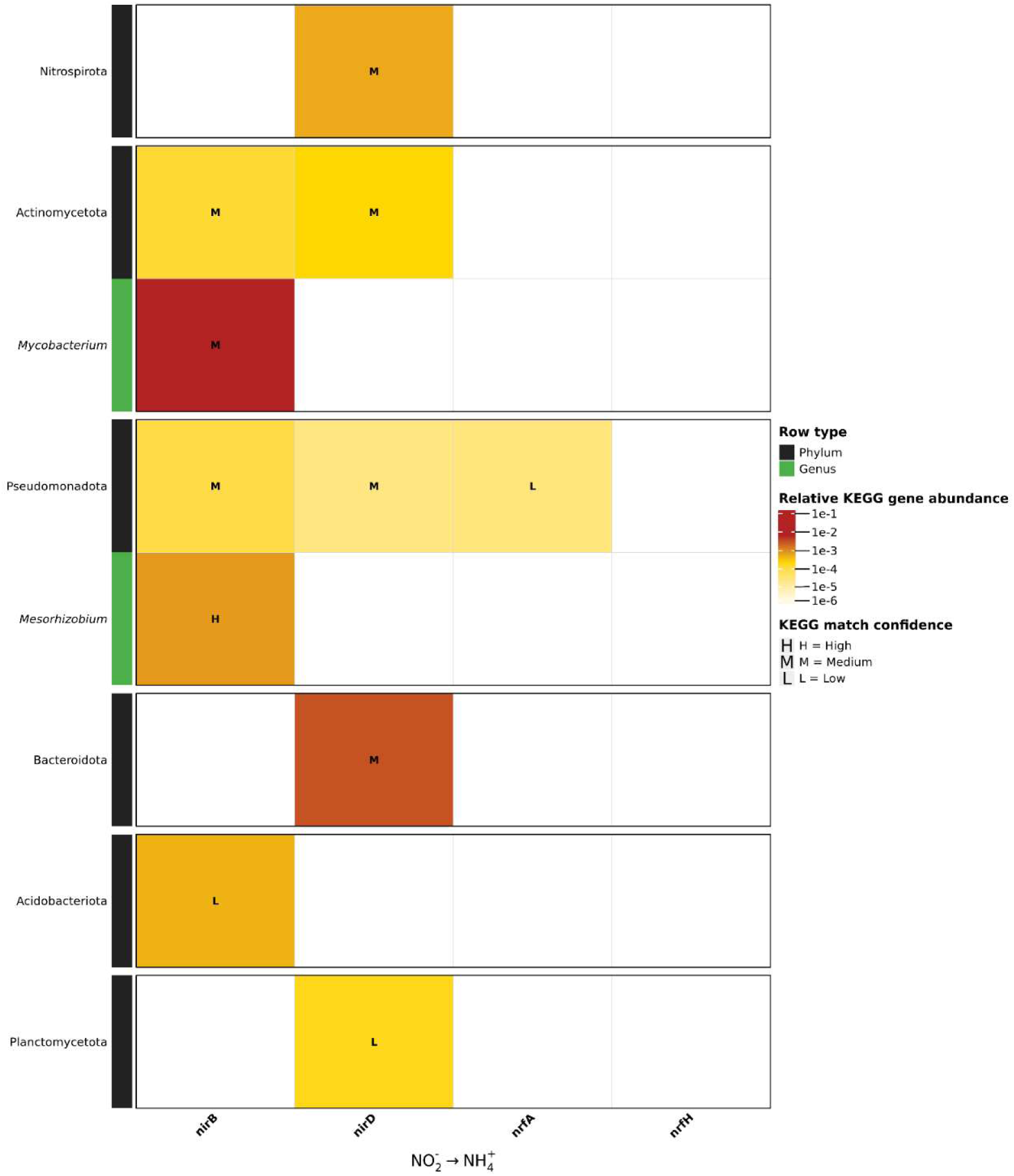
DNRA-associated genes occur across multiple lineages, including Nitrospirota. Heatmap showing the distribution of dissimilatory nitrate reduction to ammonium (DNRA) associated genes across classified phyla and genera. Rows are grouped by phylum and include phylum level summaries and resolved genus level assignments. Columns represent genes involved in nitrite reduction to ammonium. Colour intensity represents log10-normalised gene abundance, calculated from mean TPM across the three extraction replicates.

## Discussion

### Nitrogen-transformation potential was not restricted to dominant taxa

Nitrogen-transformation potential did not scale directly with taxonomic abundance. Dominant taxa, including *Nitrospiria* and Pseudomonadota, contained numerous nitrogen metabolism associated genes, whereas Actinomycetota represented only 4% of the community but included Actinomycetes, which had the highest proportion of nitrogen metabolism associated genes. Within Actinomycetota, *narGH* was assigned to *Mycobacterium* and *Mycolicibacterium*, and *nirBD* was assigned to *Mycobacterium*.

Genes associated with key pathway steps were detected in minor taxa, with *nirK* assigned to *Permianibacter* and terminal nitrous oxide reduction through *nosZ* restricted to Chloroflexota, Acidobacteriota, Myxococcota and unclassified genome bins. Monitoring dominant nitrifiers and denitrifiers alone may therefore overlook less abundant organisms encoding important pathway steps. Functional gene monitoring may complement taxonomic profiling providing information on nitrogen-transformation potential that is not detectable by taxonomic abundance alone.

### Incomplete gene systems require cautious interpretation

Several genes were detected without other subunits of their canonical complexes. Similar patterns have been reported in other SBR or related treatment environments, including *napA* expression without *napB* under SBR like conditions [99], *norB* without *norC* in wastewater treatment systems [104], and isolated *nasA/nasC* detections without *nasB* in SBR metagenomes [113, 114]. Evidence that *narGH* can retain nitrate-reduction activity without *narI* under some conditions also supports the possibility that partial systems may remain functional [95]. However, this cannot confirm activity, and some apparent partial pathways may result from incomplete assembly or genome recovery. Individual genes and missing subunits should therefore be interpreted cautiously when assessing SBR functional potential.

### Distribution of genes associated with nitrogen removal

Restricted gene distributions were identified at the beginning of nitrification and the end of denitrification. Gene-level assignments linked *amoABC* only to *Nitrosomonas* and Nitrososphaerota, indicating a narrow taxonomic distribution of ammonia oxidation potential. This restricted distribution suggests comparatively limited genetic redundancy and may make ammonia oxidation particularly vulnerable to disturbances affecting these organisms. Previous SBR studies have found that a more diverse ammonia oxidising community is more resilient to sustained loading disturbance than a less diverse nitrite oxidising community [124], and that functional gene redundancy can maintain nitrification stability [125].

A similar restricted distribution was identified at the final step of denitrification. Among classified taxa, *nosZ* was detected only in Chloroflexota, Acidobacteriota and Myxococcota, all of which were minor taxa in this SBR. Terminal nitrous oxide reduction may therefore have limited genetic redundancy and could be sensitive to community disruption. A similar relationship has been observed in other SBRs, where decreased *nosZ* abundance coincided with a strong increase in nitrous oxide accumulation [126, 127]. The restricted distribution observed here may therefore be relevant to nitrous oxide accumulation, although gene presence alone cannot establish activity or emission risk.

### Nitrogen-removal and nitrogen-retention potential

The coexistence of genes associated with denitrification, DNRA and assimilatory nitrate reduction indicates that nitrogen-removal and nitrogen-retention potential occurred within the same community. Notably, *Nitrospira* is considered a core SBR nitrifier but was also assigned *nirA*, while DNRA-associated *nirBD* genes were detected more broadly within Nitrospirota. Lineages commonly associated with nitrification may therefore also encode nitrogen-retention functions. Although gene presence does not demonstrate that these pathways were active simultaneously, these findings indicate that the functional potential of a taxonomic group cannot be inferred from its established function alone.

The broad distribution of DNRA-associated genes across multiple lineages indicates that DNRA potential is not confined to a single taxonomic group. Reactor conditions, rather than taxonomic composition alone, are likely to influence the relative contributions of nitrogen-removal and nitrogen-retention pathways. High organic carbon to nitrate ratios can favour DNRA [22], suggesting that carbon availability may influence whether nitrate is removed through denitrification or retained as ammonium through DNRA. However, longitudinal measurements of gene expression, nitrogen concentrations and operating conditions would be required before these genes can be used to define specific reactor control strategies.

## Conclusion

This study illustrates the value of gene resolved metagenomics for understanding nitrogen cycling in landfill leachate SBRs. Taxonomic abundance alone did not indicate which organisms carried the capacity for individual nitrogen pathways, or whether genes associated with nitrogen removal and nitrogen retention coexisted within the microbial community. By linking nitrogen-transformation genes to taxa and reconstructed genome bins, the analysis identified candidate low-abundance contributors, restricted taxonomic distributions of key genes, and coexisting nitrogen-removal and nitrogen-retention potential that were not apparent from taxonomic composition alone.

However, this study examined DNA-level potential in three technical DNA extraction replicates from a single sludge sample, and pathway activity cannot be established from these data. Future work linking genome resolved functional potential with expression data, reactor chemistry and nitrogen transformation rates will be needed to determine how these partitioned pathways influence nitrogen removal, retention and gas emissions in landfill leachate SBRs.

## Data availability

The raw metagenomic sequencing reads generated in this study have been deposited in the NCBI Sequence Read Archive under BioProject accession PRJNA1540796 and will be released upon publication. Analysis scripts are available at https://github.com/samuel-rattray/SBR-nitrogen-metagenomics.

## Declaration of generative AI and AI-assisted technologies in the manuscript preparation process

During the preparation and revision of this work, the authors used OpenAI ChatGPT to assist with editing analysis code and language editing, improving clarity and structure, and checking consistency within the manuscript. The authors reviewed and edited all AI-assisted output and take full responsibility for the content of the published article.

## Notes

### Competing Interest Statement

The authors have declared no competing interest.

