## Supplementary Figures for "Gene-resolved metagenomics reveals partitioned nitrogen transformation potential in a landfill leachate sequencing batch reactor microbiome"

### Supplementary Material

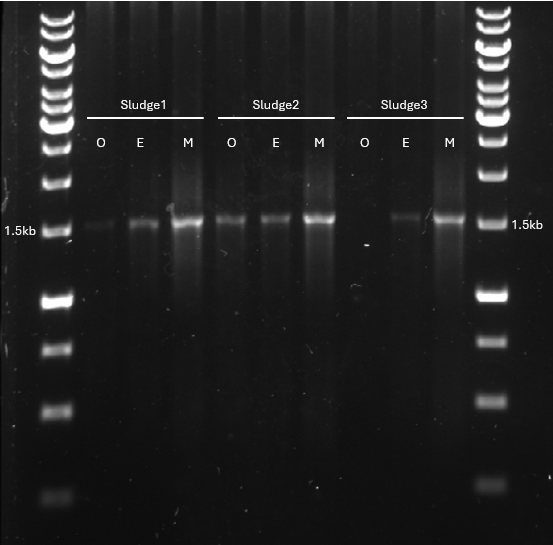

Supplementary Figure S1. Gel electrophoresis of 16S rRNA gene PCR products amplified from sludge DNA extracts before and after inhibitor removal. O, untreated extract; E, ethanol-precipitated extract; M, Monarch® PCR & DNA Cleanup Kit purified extract.

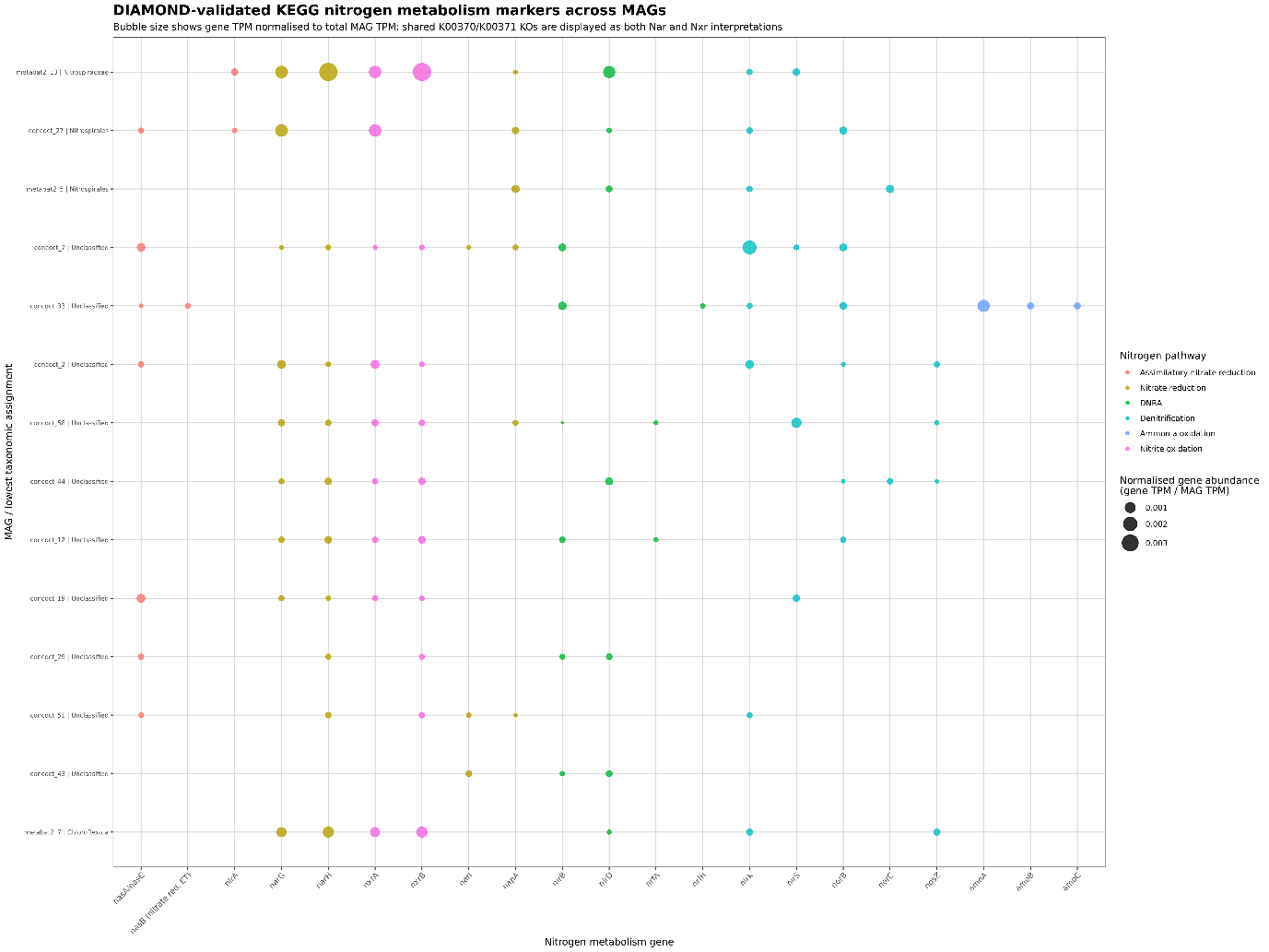

**Supplementary Figure S2. Distribution of nitrogen transformation genes across retained genome bins from landfill leachate sequencing batch reactor (SBR) sludge.** Bubble plot showing nitrogen transformation genes detected in fourteen genome bins retained at ≥30% estimated completeness and ≤10% contamination. Rows represent bins labelled by their lowest resolved taxonomic assignment and bin identifier; taxonomically unresolved bins are labelled unclassified. Columns represent nitrogen transformation genes. Bubble size indicates bin-normalised gene abundance (gene TPM divided by total bin TPM), based on mean TPM across three technical DNA extraction replicates. TPM denotes transcripts per million and is used here as a normalised metagenomic abundance measure. Colour indicates the pathway category assigned to each gene, as shown in the figure key. Bins below 50% completeness were treated as partial; missing markers do not establish biological absence. Gene detection indicates genetic potential and does not demonstrate pathway activity or completeness.

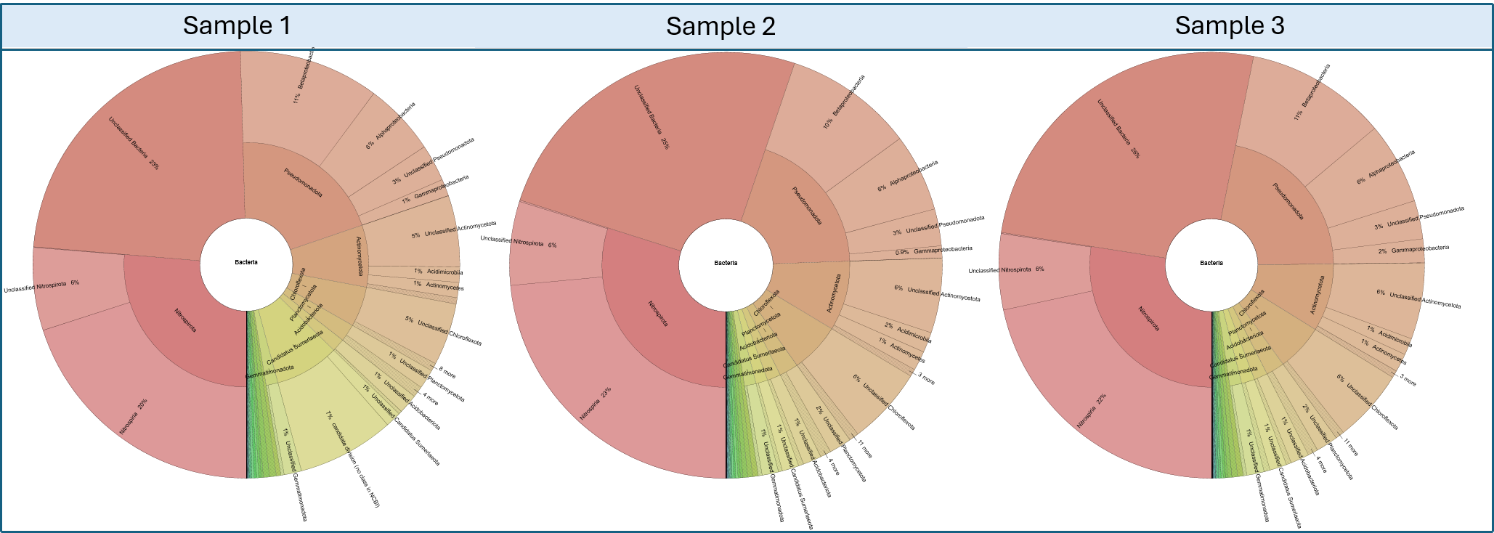

**Supplementary Figure S3. Bacterial taxonomic composition of landfill leachate SBR sludge.** Krona snapshots show all three SBR extraction replicates. Bacteria were selected as the chart root, with maximum display depth set to three. Sector proportions represent relative abundance within bacterial assignments. The complete interactive taxonomic profiles are provided in Supplementary File S1.
