## Supplementary File S1 for "Gene-resolved metagenomics reveals partitioned nitrogen transformation potential in a landfill leachate sequencing batch reactor microbiome"

Javascript must be enabled to view this page.

magnitude
magnitudeUnassigned

Sample21
Sample22
Sample23

1700332063387796

84196713541158

311

1

1

1

1

1

31

31

31

31

31

55403792560

90734

27412

27412

27412

27412

2223

3

3

3

1923

511

511

1412

1412

41119

41119

41119

41119

25216

25216

25216

25216

25216

49263362288

49263362288

49263362288

49263362288

49263362288

21

21

21

21

21

511

3

2

2

2

1

1

1

211

211

211

211

122650

1927

1927

1927

1927

102443

102443

102443

102443

1

1

1

1

1718

1116

1116

1116

1116

61

41

41

41

2

2

2

1

1

1

1

29114

29114

29114

29114

29114

1581367

47415

81

61

61

2

2

31212

31212

31212

822

822

822

1

1

1

1

98845

98845

98845

98845

1316

1316

1316

1316

3

3

1

1

1

2

1

1

1

1

1601276

1261052

435

435

435

122747

55

13

42

108735

108735

97

97

34224

34224

34224

34224

52

51

51

2

2

21

21

1

1

1

1

1

1

1

1

1

1

1

32

31

31

31

31

31

1

1

1

1

1

1

1

1

1

1

1

34324174

41

41

2

2

2

21

2

2

1

1

1491579

1491579

1491579

1491579

1491579

84435

78433

2

2

2

57

57

57

73424

73424

73424

61

61

61

61

1

1

1

1

715

715

715

715

715

99454

98254

97254

2

2

1518

1416

12

11

11

1

1

80143

80143

1

1

1

12

12

12

12

1

1

1

1

1

1

30333132

312

312

312

312

312

32220

3118

2

2

2

98

1

1

88

88

11

11

11

189

189

189

1

1

1

122

122

122

122

1611457

1611457

1611457

1611457

1611457

711132

711132

701131

59923

59923

1128

1128

11

11

11

715

715

52

52

52

213

213

213

29416

29416

12

11

1

1

1

1

1

1

1

28413

2

2

23310

23310

313

313

1

1

1

1

1

1

11

11

11

11

11

11

1

1

1

1

1

1

311

311

311

11

11

11

21

21

21

1

1

1

1

1

1

20971651006

111

11

11

11

11

1

1

1

1

1413105687

1413105687

1413105687

1413105687

1413105687

28219

28219

28219

28219

28219

45532204

27719109

27719109

27719109

27719109

60431

3

3

3

57431

197

197

38424

38424

1

1

1

1

50225

33

33

33

33216

21

21

31215

31215

146

146

146

1

1

1

1

67738

12

12

12

46323

46323

46323

53

42

42

11

11

15410

712

712

11

11

728

726

2

2002595

61

61

61

61

70934

66

1

1

65

65

56826

56826

56826

812

812

812

1221660

1221660

1221660

1221660

2

2

2

2

11

11

11

11

11

11

71

2

2

2

2

2

51

51

51

51

51

1487124741

33115

1614

1614

1614

1614

1511

1511

1110

1110

41

41

2

2

2

2

1

1

1

1

1

179

179

179

179

179

1160106581

1160106581

1160106581

1160106581

1160106581

120776

98

98

98

98

43435

24

11

11

13

13

36

36

36

38425

38425

38425

68333

68333

68333

68333

139753

56228

56228

56228

56228

77522

2226

2

2

2026

2026

141

141

141

33

22

22

11

11

38312

38312

38312

11

11

11

11

52

2

2

2

1

1

1

1

1

1

21

21

21

2

2

2

2

2

914

5

5

5

5

414

414

413

413

1

1

821

821

821

821

821

23

23

23

23

23

23

23

23

23

23

23

23

37416

412

412

412

412

412

33314

2

2

2

2

81

81

2

2

61

61

23313

23313

23313

23313

1305608718

254524186

254524186

254524186

254524186

254524186

105184532

105184532

105184532

105184532

105184532

1

1

1

1

1

1

2

2

2

2

2

2

25213

11

11

11

11

11

23112

2

2

2

2

1

1

1

1

20112

20112

96

96

1116

1116

2

2

2

2

2

29213

2128

2128

2128

2128

2128

2

2

2

2

2

65

65

65

65

65

35224

2

2

2

2

2

22120

62

42

21

21

21

21

2

2

2

719

719

719

719

2

1

1

1

1

1

1

97

56

56

56

21

21

21

2

2

2

1114

1114

1114

1114

1114

1

1

1

1

1

1

1511167

1

1

1

1

1

1501167

1491164

1461164

109850

109850

37314

37314

3

3

3

13

13

13

13

3

3

3

3

3

3

819

819

819

819

819

819

1441163

1441163

1441163

1441163

1441163

1441163

961037

961037

913

913

913

913

87934

87934

87934

87934

1501471

512

1

1

1

1

511

311

311

311

2

2

2

52

52

52

52

52

109956

109956

109956

109956

109956

1236

32

32

32

32

934

934

934

934

142

5

5

5

5

92

21

21

21

71

71

71

11

11

11

11

11

11

1

1

1

1

1

1

1

1

311

311

11

11

11

21

21

21

24

4

4

4

4

4

2

2

2

2

2

1626514428915

76891739

72

62

62

61

1

1

1

1

19911

19911

19911

1

18911

1114

51

51

51

21

21

21

32

2

2

3

3

11

11

11

96

84

32

32

52

52

12

12

12

3

3

3

3

22

11

11

11

1

1

1

1

1

1

611

31

31

31

27

27

27

2

2

2

11

1

1

1

1

16220406

11

11

11

15819398

12613331

1

4

12613326

23343

23343

722

1

622

2122

11

221

317

317

317

513

41

21

21

2

2

13

13

13

84

74

42

42

32

31

1

1

1

1

21

21

21

21

64

12

12

12

21

21

21

1

1

1

1

1

1

11

11

11

541026

531026

521025

1

612

45923

11

11

1

1

1

53331

53331

53331

53331

37739203

37739203

37739203

37739203

20312

1427

1

1

1

1

23

23

1123

1123

414

414

414

21

1

1

11

11

3

1

1

1

2

1

1

1

1

2

2

2

2

1

1

1

1

18412

18412

14310

14310

1

1

312

312

11

1

1

1

1

1

1

81807584413

33643941498

1

1

1

11

11

11

110948

1

1

101745

73533

28212

823

823

30943611372

513

1

413

26503021184

23392781044

1

29414

812

84

1

24918114

32

1214

43958185

43958185

4

1

1

3

3

2

2

2

1512476

1512476

1512476

11

11

11

18220

32

31

31

1

1

31

1

1

21

21

6110

6110

6110

617

617

617

103466925

19314

11

11

14212

714

718

31

31

11

11

3

3

3

136355

65227

65227

153

1

143

1

1

12

12

41

41

1

1

2

2

4518

4518

32

12

2

11

1

1

102456

102456

102456

664203

13

2

11

2

2

1

1

2

2

147

137

1

187

117

7

324167

324167

11

11

1

1

11

11

1

1

25

24

24

1

1

69952582

69952582

69952582

107

11

11

12

12

34

34

1

1

4

4

22

1

1

1

12

12

12

1501258

1501258

1501258

1501258

512

512

512

512

36072831908

36072831908

36072831908

36072831908

41

1

1

1

1

31

3

3

3

1

1

1

615

615

615

615

615

49523932558

2413

2413

34

34

219

157

62

33323

11

11

11

11113

412

1

42

47

47

12

1

11

1

1

1

1

1

1

1

1

2129

2129

2129

1421682

64547

33

33

1

1

32316

30315

21

2

2

29225

29225

515

211

211

23

23

11

11

1

1

1

721030

721030

721030

1811185

169981

226

51

175

2

2

2

2

34215

34215

1

1

108760

76750

3210

1

1

1

1223

1223

1223

65

65

1

1

64

12

52

103156536

102055532

1

1

125554

125554

1

1

89350478

89350478

1114

1114

1114

40614

31

31

31

37613

322

322

1514

1514

1627

1627

21

21

1

1

82863406

1

1

1

612

612

1

512

1

1

1

89

87

87

2

2

24218

12112

12112

914

31

613

32

32

2

2

2

23

23

1

13

12

1

1

2

2

57523

2

2

27416

223

25213

2717

2717

1

1

44833227

44833227

44833227

132

132

132

42

42

42

1

1

1

3

1

1

2

2

19214

13

13

1417

1417

1

1

111

111

32

2

3

1381363

1013

712

31

71734

55727

34

133

56525

56525

11

11

64422

512

512

1

1

58320

2217

1

35213

41

1

1

3

3

1

1

37214

104

104

2629

42

2227

1

1

1

1

29323138

1

1

1

41318

39318

39318

2

2

1221469

1221469

1221469

1313

11

11

6

6

63

63

47319

62

62

39317

39317

2

2

1

1

1

2219

1715

1715

1

1

21

21

11

11

12

12

212

212

212

4418

42

31

11

22

22

3814

3814

1

1

1

1

23402121232

23402121232

23402121232

23402121232

33321

33321

33321

33321

12

12

2

2

1

1

23551991199

23551991199

23551991199

23551991199

23551991199

69644

11

11

11

11

11

68643

1

1

1

1

60536

33322

33322

33322

27214

27214

27214

816

816

816

816

40531

76

1

1

1

1

66

66

66

66

88

88

88

88

88

25517

19515

11511

2

2

1

1

1059

1059

11

11

11

73

73

73

11

11

11

11

51

51

51

51

31337179

1291170

2

2

2

2

1271170

93953

93953

93953

28214

28214

28214

21

21

21

42

42

42

2013

51

51

51

51

1512

2

2

2

72

32

32

3

3

1

1

64

64

64

24

24

24

771247

771247

771247

771247

771247

851347

361122

21819

21819

21819

1533

1333

1333

2

2

49225

49225

49225

49225

211

11

11

11

11

1

1

1

1

1

1

1

1

1

1

1

1

1

1271540

1051235

1051235

1051235

1051235

1051235

2235

2235

2235

2235

2235

1276112654

23110

23110

23110

23110

23110

103094543

103094543

103094543

103094543

103094543

2051595

1114

1114

1114

1114

1941491

87734

81632

81632

11

11

52

52

83645

83645

83645

24112

24112

24112

1826

1

1

1

1

1726

21

2

2

1

1

1

1

1

1524

1524

1524

42

42

22

1

1

1

12

12

12

2

2

2

2

62

31

31

31

31

31

31

3

3

3

3

1

1

1

1

25112

715

715

715

715

715

187

187

187

187

187

21207165710577

21207165710577

21207165710577

21207165710577

21207165710577

21207165710577

54

42

2

2

2

2

4

4

4

4

1

1

1

1

1

2

2

2

1

1

1

1

21

21

21

21

21

21

36

25

12

12

12

12

13

13

13

13

11

11

11

11

11

25221187911305

90332

90332

90332

90332

90332

53494472369

53494472369

53494472369

53494472369

53494472369

73

73

73

73

73

1977514298901

73269336

73269336

73269336

73269336

1904113608565

76936263515

76936263515

76936263515

112317275000

96106054266

96106054266

1801576

1801576

1439107658

2

2

127089582

1611874

42

2

2

117750

117750

117750

2

2

2

2

2014

1

1

1

1

1

204

204

204

204

204

77855843701

46325

46325

46325

46325

46325

124882574

178

52

52

52

126

35

35

91

91

22

22

22

22

1

1

1

1

80548

25112

25112

25112

55436

12

12

1127

1127

44127

44127

11

1

1

1

1

1

1

33721

1115

1115

1115

312

2

2

111

111

1

1

19514

41

1

4

716

2

614

1

835

835

3

3

86

35

13

13

22

22

51

51

51

36211

35211

1

1

1

1

1027

1027

234

1

224

1

1

1

27126137

27126137

27126137

27126137

43

42

42

42

1

1

1

71324

132

132

132

2

2

2

1

1

1

53322

1

1

1

1

1

1

2410

107

143

2438

2438

33

22

11

2

1

1

1

1

72439312

63535269

94447

11112

82334

11

49828208

49828208

1014

112

92

33210

2

1

2027

83

2

11

11

11

61221

61221

61221

21

1

1

11

11

25220

914

914

13

13

1

1

15112

1

15111

1

1

1

1

47733852379

47733852379

47733852379

47733852379

47733852379

42131198

169

169

1

1

11

1

1

148

148

116550

116550

116550

116550

1

1

1

1

14114

14114

12

12

13112

13112

25523113

20211

20211

20211

1911877

1911877

1911877

2116

1711

1711

11

11

34

34

21

21

11

1

3

3

3

1838

1

1

1837

1414

423

19212

18212

21

1

11

3

3

1628

1628

1

1

1

129183523

59142228

59142228

59142228

59142228

113846

113846

113846

113846

58733249

1

1

1

36213

36213

36213

39826165

39826165

39826165

22

1

1

21

21

23215

1011

1011

1324

1324

128353

34

34

109

109

94134

94134

2126

1425

71

62

1

1

1

1

1

1

1

1

42

42

31

31

11

11

1

1

1

1

1

1

11

11

11

11

11

11

2

1

1

1

1

1

1

1

1

1

1

36213

33111

33111

2

2

2

31111

31111

31111

312

312

312

312

312

3

3

3

3

3

3

52

32

32

32

32

32

2

2

2

2

2

20113100

20113100

20113100

20113100

20113100

20113100

20113100

81247743978

81247743978

81247743978

81247743978

81247743978

81247743978

81247743978

772271267042415

772271267042415

772271267042415

772271267042415

772271267042415

772271267042415

772271267042415

26238138

646

646

646

646

646

646

2

2

2

2

2

2

24734131

511

511

511

511

511

24233130

33330

11

11

11

32329

43

43

27323

27322

1

13

11

2

2012793

1313

1313

1313

1882690

712

712

761335

761335

1051253

90736

15517

837

837

837

837

2

1

1

1

1

1

1

1

1

1

1

2

2

2

2

2

2

2

2

2

2

2

2

11

11

11

11

11

11

21

21

21

11

11

11

11

1

1

1

1

2136

1

1

1

1

1

1

2

2

2

1

1

1

1

1

1

1

1

1

1

1

1

411

411

411

411

411

411

3

3

3

3

3

3

1

1

1

1

1

1

2

1

1

1

1

1

1

1

1

1

1

121

121

121

121

121

1

111

21

11

11

11

11

11

1

1

1

1

1

3

3

2

2

2

2

1

1

1

1

3

3

3

3

3

3

1

1

1

1

1

1
